# When DL-Based Prescreening Meets Synthon-Based Docking: Target-Adapting PharmacoNet via MEL-Steered Correction

**DOI:** 10.64898/2026.09.02.748684

**Authors:** Wenjin Liu, Yongchan Hong, Thomas Ku, Woojin Lee, Emily Nguyen, Ao Xu, Vsevolod Katritch

## Abstract

As chemical libraries expand into the trillions of molecules, Virtual SYNthon Hierarchical Enumeration Screening (V-SYNTHES) has emerged as a leading strategy for making gigascale virtual screening computationally tractable. In V-SYNTHES, a Minimal Enumeration Library (MEL) of chemical fragments is docked against a target first, and only the top-scoring fragments are expanded into full ligands for large-scale docking. However, among the large number of comparably well-docked fragments, only a small fraction can be expanded under a fixed docking budget, leaving most similarly promising fragments unexplored. General-purpose prescreening tools can be adopted to address this constraint, reallocating the same docking budget across a larger pool of fragments’ enumerated full ligands by their proxy score. However, such tools are applied without accounting for target-specific pocket environments. One such method, PharmacoNet, predicts interaction hotspots from a protein structure and ranks candidates via graph matching against a fixed set of interaction-type weights. We recognize that V-SYNTHES’s initial fragment-docking step, ordinarily used only for selection of best fragments for expansion, already reveals which of these hotspots and interaction types a given pocket actually favors, and we can recover this signal to fine-tune PharmacoNet accordingly. We introduce MEL-Steered PharmacoNet, a parameter-efficient adaptation framework that specializes PharmacoNet to a given target through two composable mechanisms: (i) empirical density-map steering of predicted pharmacophore hotspots, and (ii) empirical fine-tuning of interaction-type scoring weights. Across three structurally distinct GPCR targets (CB2, GPR91, 5-HT_2*A*_R), MEL-Steered PharmacoNet achieves substantial enrichment factor (EF_100_) gains over a random baseline, and improves EF_100_ over PharmacoNet by 8.94×, 6.87×, and 1.69×, respectively. The fitted per-target weights further reveal distinct, chemically interpretable interaction profiles that PharmacoNet’s generic fixed weights fail to capture. These results show that fragment-docking data already generated by the standard V-SYNTHES pipeline can adapt a general-purpose pharmacophore prescreening method to an individual target, significantly improving its performance while retaining its ultra-fast screening ability, with no additional experimental data or model retraining.

## 1. Introduction

Modern drug discovery increasingly draws on chemical libraries that have expanded into the trillions of molecules, and structure-based virtual screening is central to navigating such chemical spaces. However, its docking step remains prohibitively expensive to apply exhaustively across libraries of this scale, motivating a broad class of fast prescreening methods that prioritize candidates for downstream docking [4, 5]. Pharmacophore-based prescreening is one such approach: PharmacoNet, for instance, uses deep learning to construct an explicit three-dimensional pharmacophore model based on a target pocket and ranks candidates by geometric complementarity to that model [11].

Synthon-based hierarchical enumeration screening, exemplified by V-SYNTHES (Virtual SYNthon Hierarchical Enumeration Screening) [9] (§2.2), is a major approach to drug discovery focused explicitly on combinatorial chemical spaces. These spaces are organized around a fixed set of reactions, each defining a scaffold with multiple attachment points that building-block synthons can be combined into to assemble a compound. V-SYNTHES explores such spaces far more efficiently than exhaustive screening by exploiting their inherent hierarchical structure: rather than docking fully elaborated compounds directly, it first docks a small library of chemical fragments against the target, and only the top-scoring fragments are then combinatorially expanded into fully enumerated compounds for large-scale docking. Enumerating promising fragments into full ligands yields a substantially stronger candidate pool than an equivalently sized random set of ligands drawn from the full library. For instance, V-SYNTHES workflow alone achieves an EF_100_ of 1158.6× over a random baseline of the same size on CB2 (Table 2).

However, at the scale of these combinatorial chemical spaces, a large number of fragments achieve comparably strong docking scores, such that the top of the ranking is effectively flat and offers little to discriminate among them. This flatness directly constrains the fixed docking budget. Expanding fragments into fully enumerated compounds and docking them in strict rank order, following the V-SYNTHES workflow, exhausts the 1M-compound budget after only the top 82, 88, and 97 fragments for CB2, GPR91, and 5-HT_2A_R, respectively. The vast majority of similarly well-docked fragments, and the compounds enumerated from them, therefore go unexplored. General-purpose prescreening tools are often applied to address this constraint, enabling practitioners to enumerate a larger set of similarly well-docked fragments into full ligands, reallocate the fixed docking budget across this larger pool, and select which compounds are most worth docking regardless of their original fragment rank.

On three GPCR targets under a common docking protocol, Table 2 compares the enrichment achieved by V-SYNTHES’s strict fragment-rank-order selection against PharmacoNet’s proxy-score selection, each independently shortlisting one million compounds to dock from the same enumerated pool. On two of the three targets, PharmacoNet yields lower enrichment than V-SYNTHES; on CB2, EF_100_ falls from 1158.6× to 225.3×, a more than fivefold reduction. On the remaining target, it yields enrichment comparable to V-SYNTHES rather than a clear improvement. A related pattern holds for other generalist prescreening methods: DrugCLIP [3] and LigUnity [2] likewise reduce enrichment significantly relative to V-SYNTHES on two of the three targets, though on the third (GPR91) both instead yield an improvement over V-SYNTHES (Table 2).

This inconsistency does not indicate a deficiency in PharmacoNet’s underlying pharmacophore model. Rather, it reflects a direct consequence of the manner in which such models are trained and deployed: PharmacoNet is trained on PDBbind v2020 [7], a fixed set of roughly 17,600 crystallized protein–ligand complexes, and applied unmodified to every target pocket it subsequently encounters, with no mechanism to adapt to target-specific preferences. In practice, the benefit of prescreening therefore largely depends on how closely a given target’s interaction profile resembles the population PharmacoNet was trained on, a correspondence that cannot be known in advance.

The V-SYNTHES workflow offers insights into the signal needed to resolve this limitation, as a routine byproduct. The first step of V-SYNTHES docks the fragment library against the protein target to rank fragments (§2.2). This docking result also reveals, empirically, which interaction types and geometries that a protein pocket favors. Typically, this signal is discarded once the top-scoring fragments are selected for subsequent expansion. To our knowledge, we are the first to recover this discarded signal and use it to adapt a general-purpose prescreening method to an individual target. Our main contributions are as follows:

- We identify the fragment-docking step of the standard V-SYNTHES workflow as an existing, discarded source of target-specific signal, and show that this signal, rather than additional experimental data or retraining on a larger complex database, is sufficient to convert a general-purpose prescreening method into a target-specific one that reliably improves screening performance while retaining its ultra-fast screening ability (Supplementary Info §S1).
- We propose MEL-Steered PharmacoNet, which specializes PharmacoNet to a given target through two composable corrections based on empirical fragment-docking data: geometric refinement of its predicted pharmacophore hotspots, and calibration of its interaction-type scoring weights.
- Across all three benchmarked targets and both enrichment metrics (EF_100_ and EF_1000_), MEL-Steered PharmacoNet outperforms both PharmacoNet and the V-SYNTHES baseline. As a further contribution of the weight correction, our method yields an interpretable, pocket-specific set of scoring weights across interaction types for each target, revealing chemically distinct interaction profiles that PharmacoNet’s fixed, heuristically-assigned weights cannot capture.

## 2. Related Work

### 2.1 Virtual Screening

Virtual screening (VS) sits at the hit discovery stage of drug discovery, after target identification and before hit-to-lead and lead optimization. Given a target, VS computationally ranks candidates from a large compound library by predicted binding likelihood — via structure-based methods (e.g., docking [14], pharmacophore matching [17]) or ligand-based methods (e.g., QSAR [13], ML scoring [12]) — to narrow the library down to a manageable shortlist for wet-lab testing. As standard practice at this stage, VS methods are evaluated entirely on computational proxy scores such as docking scores. Experimental binding data is costly and low-throughput, so it is collected only once a shortlist has already been selected at later stages; it is therefore not used to evaluate VS methods themselves.

### 2.2 V-SYNTHES

V-SYNTHES [9] established synthon-hierarchical enumeration as a practical alternative to brute-force docking of ultra-large libraries: docking a small representative fragment library first, expanding only the top-scoring fragments into full compounds, and docking those compounds in turn. By strategically docking a small fraction of combinatorial libraries, V-SYNTHES reported over a multi-thousand-fold reduction in docking cost while still recovering top-scoring compounds, and has since been widely adopted as a standard approach for gigascale structure-based screening.

### 2.3 PharmacoNet

PharmacoNet [11] is the pharmacophore-based prescreening method our approach builds on and corrects, and we treat it here as our base method. Given a protein pocket structure, a deep learning model predicts a set of interaction hotspots (locations, together with an interaction type and confidence, where a ligand functional group would need to sit to form a favorable non-covalent interaction) without reference to any candidate ligand. A candidate ligand is then scored by a coarse-grained graph-matching procedure [11] that aligns its own functional groups against these hotspots, using a fixed set of seven per-type scoring weights and a lightweight analytical function to evaluate geometric fit. This scoring machinery is established once across a broad population of protein–ligand interactions and then applied unmodified to every pocket it encounters, with no mechanism to incorporate target-specific evidence.

### 2.4 Embedding-Based Prescreening Methods

DrugCLIP [3] and LigUnity [2], two embedding-based generalist prescreening methods that share the same target-agnostic limitation as PharmacoNet, are described in Supplementary Info §S2.

## 3. Preliminaries

### 3.1 PharmacoNet Scoring Formalism

The PharmacoNet scoring formula operates over two node sets: the pharmacophore model and the candidate conformer. The pharmacophore model is a set of model nodes predicted hotspots, each with an interaction type, center, and extent. The candidate conformer is decomposed by functional-group typing into a parallel set of ligand nodes. Graph-matching assigns ligand nodes to model nodes, a single matched pair being one ligand node i mapped to one model node j . Scoring is computed on edges, each joining two matched pairs (i → j) and (i^′^→ j ) and comparing an observed ligand-side length against a predicted model-side distribution:

- Ligand-side (observed): 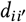, the distance between ligand nodes i and i^′^.
- Model-side (predicted): 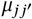, the center-to-center distance between j and j ^′^; 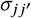, the average of their radii; and per-type weights W_t ( j )_, 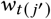 .

Each edge contributes a distance likelihood evaluating the observed ligand separation against the model-predicted distribution,

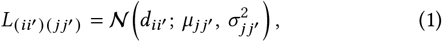

scaled by the two model nodes’ per-type weights 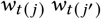. The total score S sums these over all edges,

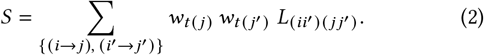

For clarity, equations 1–2 present a simplified single-Gaussian form. The full form, in which L is a Gaussian mixture and the weights enter both the mixture coefficients and the pair weight, is given in [11].

Graph-matching selects the assignment maximizing S under geometric compatibility constraints. As each term depends on the weights only through the positive product 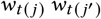, scaling the per-type weights by a common positive constant rescales S identically across assignments, leaving each candidate’s best assignment and the overall ranking unchanged. Only the ratios between the seven weights, not their absolute magnitudes, therefore determine which interactions dominate a match and how candidates rank relative to one another.

### 3.2 Minimal Enumeration Library

A Minimal Enumeration Library (MEL) fragment consists of a chemical scaffold, which persists unchanged into the final molecule, and one or more placeholder cap groups at the remaining attachment point(s), which are later replaced by real synthons during full enumeration. Because docking every fully enumerated compound in a trillion-sized combinatorial library is computationally prohibitive, MEL construction instead enumerates only one R-group position per reaction, capping the remaining position with a minimal placeholder isotoped for identification, methyl or phenyl depending on the reaction chemistry at that position, yielding a compact representative set that covers the reaction’s scaffold diversity at a fraction of the docking cost. Docking this Minimal Enumeration Library, rather than the full compound space, is what makes both initial fragment prioritization (§1) and the target-specific correction introduced in the sections that follow computationally feasible.

## 4. Methodology: MEL-Steered PharmacoNet

Our MEL-Steered PharmacoNet methodology is shown in Figure 1^2^. Code and data used in this work will be released publicly upon publication.

**Figure 1.**
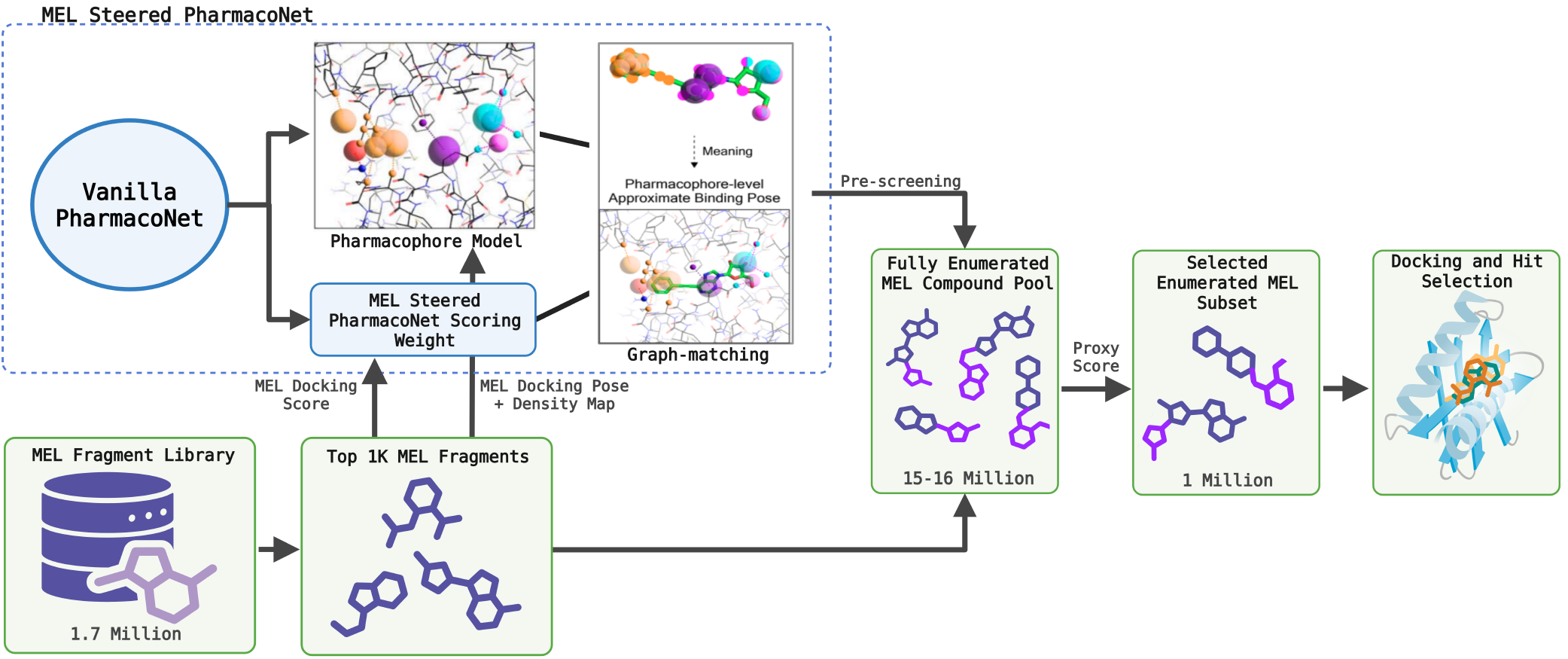
Overview of MEL-Steered PharmacoNet. Docking the MEL fragment library against the protein target both ranks fragments for enumeration and yields empirical evidence that corrects PharmacoNet’s predicted density fields and interactiontype weights; the corrected model then prescreens the enumerated compound pool.^1^

### 4.1 MEL-Steered Geometry Refinement

#### Empirical density construction

For each target, the MEL fragment library is docked against the protein pocket, and the top-scoring fragment docking poses are retained (§6). We run the Protein– Ligand Interaction Profiler (PLIP) [10] on each fragment–protein complex to identify confirmed, geometrically realized interactions, recording the ligand-side coordinate and interaction type of each. For each interaction type, PLIP contacts are partitioned by their structural origin (§3.2): a real-synthon field generated from scaffold-atom contacts, and a cap-only field generated from cap-group contacts. Within each field, every confirmed contact of a given interaction type contributes an isotropic three-dimensional Gaussian centered on its voxel position, and contacts of the same type are combined by summation, so a location supported by contacts from several different fragments accumulates a stronger signal than one supported by a single fragment. The summed grid for each type is then max-normalized to [0, 1].

#### Augmentation

The empirical fields correct PharmacoNet’s deep-learning-predicted density rather than replacing it, preserving prior coverage where the fragment data does not sample. We evaluate full replacement as an ablation in §6.4.

Within each field, candidate sites are the connected components of the surviving signal, following PharmacoNet’s convention that each segment of a thresholded density map is a distinct pharmacophore point. Let D_d,h,W_ ∈ [0, 1] be the predicted density at grid index [ d, h, W], M the voxel indices of one segment with | M | their count, and T : ℤ^3^ → ℝ^3^ the map from grid index to spatial coordinate. Each site is summarized by

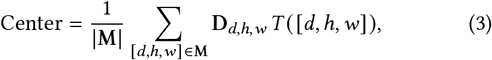

weighting each voxel’s coordinate by its density and normalizing by the voxel count. Sites are matched to same-type deep-learning hotspots by Euclidean distance between these centers, accepted at a 2.0 Å threshold. Combination is applied per site according to source:

- Real-synthon sites, with a matched hotspot:

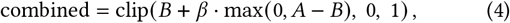

restricted to B’s footprint, where B is the empirical field, A the matched hotspot’s density, and β an agreement-bonus coefficient. Without a match, the site is retained unmodified.
- Cap-group sites, with a matched hotspot:

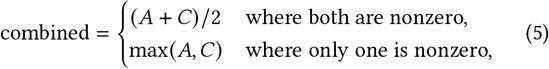

where C is the cap-only empirical field, evaluated over the full grid rather than restricted to the site’s footprint. Without a match, the site is discarded.
- Unmatched deep-learning hotspots are retained individually at reduced confidence.

This produces one combined density field per interaction type, integrating empirical and deep-learning-predicted signals according to the rules above.

#### Clustering

We convert each combined field into graph nodes following PharmacoNet’s two-stage clustering procedure. Prior to clustering, voxels are thresholded according to the source of their signal: voxels carrying empirical signal are retained since that signal originates from a confirmed interaction rather than an uncertain prediction, while voxels carrying only deep-learning-predicted signals are thresholded at 0.5, PharmacoNet’s default cutoff. In the first stage, surviving voxels are grouped into connected components under 26-connectivity (any two voxels sharing a face, edge, or corner are treated as connected), and components smaller than a minimum voxel count are discarded; each surviving component with binary mask M becomes a node, with center given by the intensity-weighted centroid of its voxels (Eq. 3) and radius back-calculated from voxel count under a spherical-volume assumption (Eq. 6),

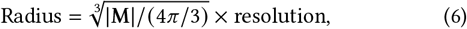

where resolution is the grid spacing. In the second stage, nodes belonging to a “major” interaction type (cationic, anionic, and aromatic-forming types) absorb any nearby node of a “minor” type (e.g., hydrogen-bonding, hydrophobic) within a fixed proximity radius of 3.0 Å, forming the clusters that serve as the model nodes in graph-matching (§3.1). Nodes without a nearby major-type partner are instead grouped with other same-type nodes within that same radius. Graph-matching aligns ligand functional groups to these nodes by comparing the ligand’s internal interatomic distances against the corresponding node-to-node distances. Applying this clustering procedure to the post-augmentation field yields a single pharmacophore model in which hotspot geometry is corrected by empirical fragment evidence wherever such evidence is available, and governed by the original deep-learning prediction elsewhere.

### 4.2 MEL-Steered Weight Calibration

#### Reparametrization

Weight calibration is fit on top of whichever pharmacophore geometry is already in place and is performed independently of that choice, making the two corrections sequential rather than competing. By the ratio-invariance property established in §3.1, a uniform rescaling of all seven interaction-type weights (Supplementary Info §S4) leaves every score comparison unchanged: we therefore fix W_hydrophobic_ = 1.0 as an anchor and calibrate the remaining six weights (W_aromatic_, W_HBD_, W_HBA_, W_halogen_, W_cation_, W_anion_) relative to it, searched log-uniformly over [0.5, 24.0].

#### Objective

Calibration is performed per pocket, against that target’s curated top-scoring MEL fragment population — the same population used to construct the empirical density fields in §4.1. Each fragment’s docking score is converted to a quality label y = −(ICM score), so that higher values indicate better binders, matching PharmacoNet’s own score convention. Writing ŝ (W) for PharmacoNet’s predicted scores under weight vector W and y for the docking-score-derived labels, the six free weights are selected to maximize:

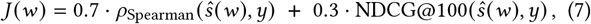

where NDCG is the Normalized Discounted Cumulative Gain [15]. The Spearman term rewards global rank agreement, while NDCG@100 concentrates fitting pressure on correctly ordering the top of the ranked list, which is what enrichment-based screening ultimately rewards. Because J is calculated through a non-differentiable graph-matching procedure, gradient-based optimization is not applicable; we instead use Optuna’s Tree-structured Parzen Estimator [16], a Bayesian, sequential model-based black-box optimizer, run for 1000 trials per study. Weights are selected via 5-fold cross-validation and refit on the full population for deployment (Supplementary Info §S3); the resulting, independently-fit weight vector for each target is referred to as MEL-Derived ScreenWeights throughout the remainder of the paper.

## 5. MEL-Signal Validation

Before evaluating downstream screening performance, we verify that the empirical signal used by the geometry refinement and weight calibration behaves as intended: §5.1 examines the geometry refinement (§4.1), and §5.2 the weight calibration (§4.2).

### 5.1 Geometry-Correction Sanity Checks

To verify that the geometry refinement responds differently depending on whether empirical evidence agrees with, partially agrees with, or contradicts the deep-learning prior, we compare PharmacoNet’s predicted density map, the empirical density map, and the resulting MEL-steered density map across representative interaction types and targets. For GPR91, as shown in Figure 2(a),

**Figure 2.**
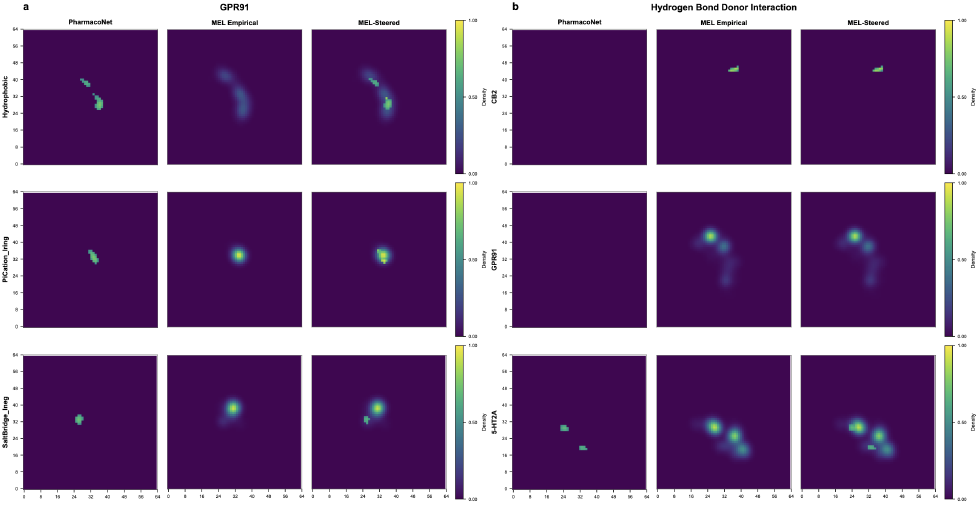
Comparison of PharmacoNet’s predicted density map, the empirical density map, and the resulting MEL-steered density map. (a) Representative interaction types on GPR91 — hydrophobic contact, cation–π, and salt-bridge. (b) Hydrogen-bond-donor interaction density maps across CB2, GPR91, and 5-HT_2*A*_R.

PharmacoNet predicts two spatially concentrated hotspots for hydrophobic contacts, while the empirical map instead shows broad, diffuse regions of moderate density with no comparably sharp peak; the deep-learning prediction falls within the empirical footprint, indicating agreement on location but a substantial difference in spatial concentration and signal magnitude. The steered map preserves the empirical map’s broad extent while intensifying the sub-region overlapping PharmacoNet’s prediction. For the cation– π (PiCation_lring) interaction, PharmacoNet and the empirical map instead agree closely in both location and intensity, and the steered map correspondingly shows a single, reinforced hotspot at the shared location. For the salt-bridge (SaltBridge_lneg) interaction, PharmacoNet and the empirical map disagree in location entirely, with a Euclidean distance between their weighted centroids exceeding the 2.0 Å agreement threshold used for matching (§4.1). Here, the steered map retains both as separate hotspots, reflecting §4.1’s rule that unmatched empirical and deep-learning evidence are preserved independently. Figure 2(b) extends this comparison across targets, examining the hydrogen-bond-donor (HBond_ldon) interaction on CB2, GPR91, and 5-HT_2A_R. On CB2 and GPR91, PharmacoNet predicts no density for this interaction type at all, despite the empirical map confirming its presence at both targets. With no deep-learning hotspot to match against, the steered map at these two targets is determined entirely by the empirical signal, so the correction recovers interaction sites the deep-learning prior missed. On 5-HT_2A_R, PharmacoNet does predict this interaction, and two of its predicted hotspots align with two of the three empirical hotspots. In the steered map, the two agreeing hotspots are reinforced, while the third is added from empirical evidence alone, matching the same no-match preservation behavior observed for the missed interactions on the other two targets.

### 5.2 Weight-Fitting Rank Correlation

We next report how well the calibrated weights recover the fragment-docking ranking they are fit against, prior to any downstream screening evaluation. Table 1 reports, for each target, the default PharmacoNet weights’ and the optimized weights’ agreement with the MEL fragment population’s docking-score ranking, under both components of the fitting objective (§4.2): Spearman rank correlation and NDCG@100. Across all three targets, optimized weights improve both metrics over the default weights, with Spearman gains ranging from roughly 0.05 to 0.08 and NDCG@100 gains ranging from roughly 0.04 to 0.07. However, even after calibration, Spearman correlation between PharmacoNet’s predicted score and MEL docking rank remains modest. The actual measure of whether this signal translates into improved compound selection is the enrichment factor evaluation in §6.3, evaluated on each target’s fully enumerated compound library rather than on the fitting population itself.

**Table 1.** Spearman rank correlation and NDCG@100 between predicted score and MEL fragment docking rank, for default and MEL-Derived ScreenWeights across three GPCR targets.

| Target | Weights | Spearman↑ | NDCG@100↑ | Objective↑ |
| --- | --- | --- | --- | --- |
| CB2 | Default | 0.186 | 0.659 | 0.328 |
|  | Optimized | 0.236 | 0.711 | 0.379 |
| GPR91 | Default | 0.089 | 0.563 | 0.231 |
|  | Optimized | 0.164 | 0.637 | 0.306 |
| 5-HT<sub>2A</sub>R | Default | 0.093 | 0.606 | 0.247 |
|  | Optimized | 0.142 | 0.645 | 0.293 |

## 6. Experiments & Results

### 6.1 Targets & Datasets

#### REAL Space and MEL construction

The Enamine REAL Space is a make-on-demand virtual compound library comprising molecules synthesizable via optimized parallel synthesis protocols, with a synthesis success rate greater than 80% and a turnaround time of approximately four weeks [6]. The library is organized on a combinatorial, modular principle: each validated chemical reaction defines a scaffold template encoded in Markush format, with two or more attachment points (R-groups) where building-block reagents (synthons) are incorporated to assemble the final compound [6]. The version of REAL Space used in this work contains approximately 36 billion compounds built from 164 unique chemical reactions and 112,514 reactants [8]. We instantiate the MEL construction described in §3.2 on the two-component reaction subset of REAL Space, yielding approximately 1.7 million MEL fragments that collectively cover the full synthon–scaffold diversity of the two-component REAL Space.

#### Target-specific compound space

We evaluate on three structurally distinct GPCR targets: the cannabinoid receptor CB2 (PDB 5ZTY), the hydroxycarboxylic acid receptor GPR91 (PDB 6RNK), and the serotonin receptor 5-HT_2A_R (PDB 7WC6). For each target, the MEL fragment library is docked against the target using ICM-Pro, and the top-1,000-scoring fragments are retained and combinatorially expanded (each fragment’s capped position replaced by its full range of compatible synthons) into a candidate compound set specific to that target, filtered for physicochemical and substructure quality (Supplementary Info §S5). The result is three target-specific, ∼15-million-compound enumerated pools, forming the common candidate space over which every method is evaluated in what follows.

#### Target-agnostic decoy set

To evaluate enrichment, we construct a target-agnostic decoy set of one million random compounds, shared across all three targets and all methods compared. For each MEL, we randomly select 1% of the available synthons at its Rgroup positions, yielding 6,418 synthons in total across all two-component reactions, and all combinatorially possible products of these synthon subsets are enumerated. The resulting pool is filtered by Lipinski’s rule of five, and one million compounds are drawn at random from the filtered pool to form the decoy set. This same set of one million compounds is then docked independently against each of the three receptor structures, providing a target-specific baseline for the enrichment achievable by docking an equal-size set of randomly selected compounds.

### 6.2 Baselines & Evaluation Protocol

#### Baselines

We compare MEL-Steered PharmacoNet against four methods. V-SYNTHES selects which compounds to dock by expanding top-docking fragments in rank order, under the fixed docking budget of one million compounds. PharmacoNet is evaluated as a prescreening method that instead prioritizes docking by its own predicted score over the same enumerated compound space, using its unmodified, population-trained pharmacophore geometry and default scoring weights. DrugCLIP and LigUnity are evaluated identically, as embedding-based prescreening methods over the same enumerated space; as discussed in Supplementary Info §S2, both are trained and validated primarily against experimental binding affinity rather than docking score, but we include them nonetheless since embedding-based prescreening of this kind remains common practice among practitioners. For every prescreening method, including MEL-Steered PharmacoNet, the top one million compounds by that method’s native score are selected and docked, holding the docking budget fixed at 1 million across methods to ensure a fair comparison.

#### Docking protocol

All compounds are docked using ICM-Pro v3.9-3b (Molsoft LLC) [1] under an identical protocol across all methods and targets. Docking samples the ligand’s internal coordinates within a rectangular box centered on the orthosteric binding pocket, using biased-probability Monte Carlo (BPMC) optimization of the ligand’s internal coordinates against precalculated grid energy potentials of the receptor, at a thoroughness parameter of 2. Following field convention, each compound is docked once: BPMC’s internal search already samples multiple starting orientations per run and reports the best-scoring pose as the docked pose, so a single run approximates what replicate docking would otherwise provide. Docking produces a Dock Score, for which more negative values indicate stronger predicted binding.

#### Evaluation metrics

Enrichment factors were used to evaluate how effectively each screening method concentrates high-scoring compounds relative to a random baseline of equal size, under identical docking conditions. All enrichment calculations are purely docking-score based and do not involve experimental binding data. The active set for each method is its corresponding selected pool of approximately one million fully enumerated compounds, described above. We report enrichment factors at two operating points, EF_100_ and EF_1000_, computed by sweeping a docking-score threshold down the ranked active list: at each threshold, the number of actives (N_active_) and decoys (N_decoy_) scoring at or better than it is recorded, and the enrichment factor is computed as

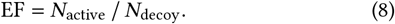

An exponential curve is fit to the resulting enrichment values to reduce noise from small active counts (*<* 50) at the top of the ranking; EF_100_ and EF_1000_ are then read off this fitted curve at the thresholds where the active count first reaches 100 and 1,000, respectively.

### 6.3 Main Results

Table 2 and Figure 3 report EF_100_ and EF_1000_ for all baselines and MEL-Steered PharmacoNet, across all three targets. Every benchmarked method achieves large EF values: on CB2 alone, EF_100_ ranges from 37.2× for DrugCLIP to 2013.8× for MEL-Steered PharmacoNet. This is expected: all of them screen the fragment-curated, target-specific compound pool, which is far more enriched than an equally sized random set, as it was built by exploiting the hierarchical structure of combinatorial chemical space (§1 & §6.1). Each method, however, is permitted to dock only one million compounds from this pool (§6.1), and MEL-Steered PharmacoNet allocates that budget best, improving EF_100_ over vanilla PharmacoNet by 8.94× on CB2, 6.87× on GPR91, and 1.69× on 5-HT_2A_R, with EF_1000_ gains following a similar pattern. Furthermore, it exceeds the high-performing, widely used V-SYNTHES workflow on all three targets: a 74% relative gain in EF_100_ on CB2, a 2.98-fold gain on GPR91, and a 75% gain on 5-HT_2A_R. Among all prescreening methods evaluated, MEL-Steered PharmacoNet is the only one that consistently improves on this already-efficient workflow rather than degrading it on some targets, as vanilla PharmacoNet does (Table 2). Against the two learned-embedding baselines, DrugCLIP and LigUnity, the margin is comparably decisive; LigUnity’s EF_100_ on GPR91 (595.3× ) is the closest any baseline comes to MEL-Steered PharmacoNet’s performance (774.8× ) across the full table, and remains behind on both metrics at that target as well as at the other two.

**Figure 3.**
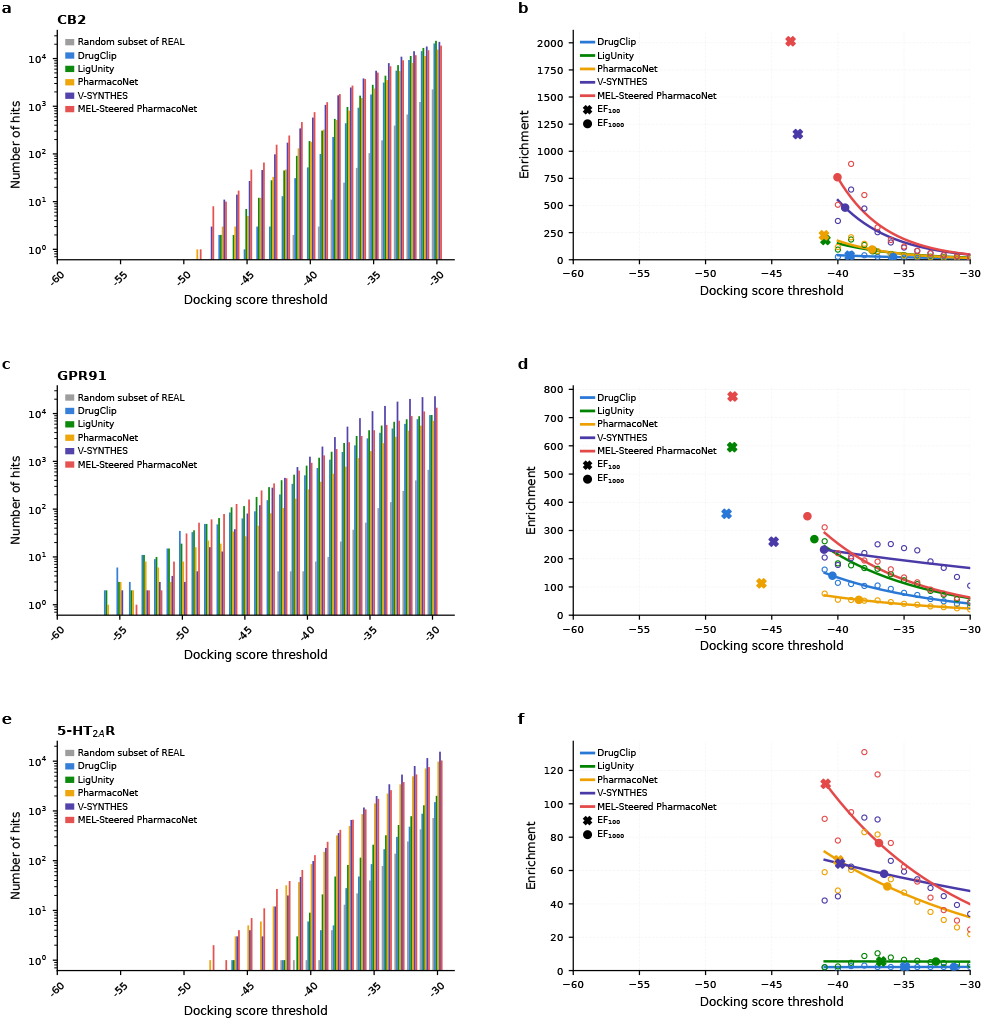
Assessment of MEL-Steered PharmacoNet performance relative to V-SYNTHES and other pre-screening methods across three receptor targets.

**Table 2.**
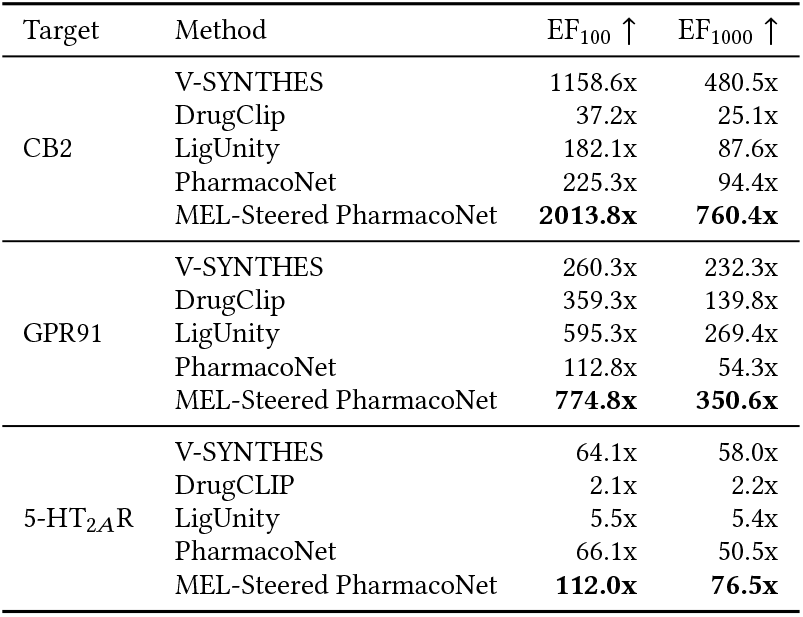
Enrichment factors (EF_100_, EF_1000_) for all baselines and MEL-Steered PharmacoNet, across three targets.

### 6.4 Ablation Studies

To isolate the individual contribution of each correction, Table 3 reports the full five-row ladder from vanilla PharmacoNet through each single correction to the fully combined method, across all three targets.

**Table 3.** Ablation: EF_100_ and EF_1000_ for vanilla PharmacoNet, each single correction, and the fully combined method, across three targets.

| Target | Method | EF<sub>100</sub> ↑ | EF<sub>1000</sub> ↑ |
| --- | --- | --- | --- |
| CB2 | PharmacoNet | 225.3x | 94.4x |
|  | PharmacoNet + Empirical ScreenWeights | 344.9x | 136.7x |
|  | PharmacoNet + Empirical-Only GeometryRefine | 856.6x | 324.7x |
|  | PharmacoNet + DL&Empirical GeometryRefine | 907.5x | 345.2x |
|  | MEL-Steered PharmacoNet | 2013.8x | 760.4x |
| GPR91 | PharmacoNet | 112.8x | 54.3x |
|  | PharmacoNet + Empirical ScreenWeights | 646.9x | 222.3x |
|  | PharmacoNet + Empirical-Only GeometryRefine | 236.1x | 133.2x |
|  | PharmacoNet + DL&Empirical GeometryRefine | 245.6x | 137.6x |
|  | MEL-Steered PharmacoNet | 774.8x | 350.6x |
| 5-HT<sub>2A</sub>R | PharmacoNet | 66.1x | 50.5x |
|  | PharmacoNet + Empirical ScreenWeights | 83.4x | 61.2x |
|  | PharmacoNet + Empirical-Only GeometryRefine | 63.5x | 46.6x |
|  | PharmacoNet + DL&Empirical GeometryRefine | 82.6x | 58.9x |
|  | MEL-Steered PharmacoNet | 112.0x | 76.5x |

Both corrections independently improve over vanilla PharmacoNet, and combining them yields a further gain beyond either correction alone at every target and both metrics. Geometry refinement alone produces the larger single-correction gain on CB2, raising EF_100_ from 225.3× to 907.5×, while weight calibration alone produces the larger single-correction gain on GPR91, raising EF_100_ from 112.8× to 646.9×, a nearly 6-fold improvement. On 5-HT_2A_R, the two corrections are comparable in isolation (82.6× and 83.4× EF_100_, respectively) but neither alone reaches the combined method’s 112.0× . EF_1000_ follows a closely similar pattern to EF_100_ across all three targets and both single corrections (Table 3). This pattern is consistent with geometry refinement and weight calibration improving different aspects of PharmacoNet rather than one subsuming the other.

Between the two geometry-only variants, the “DL&Empirical GeometryRefine” variant uses both PharmacoNet’s DL-predicted signal and the MEL-derived empirical signal and outperforms the empirical-only variant at every target. This advantage of retaining both signals is most pronounced on 5-HT_2A_R (82.6× vs. 63.5× EF_100_), consistent with the design rationale in §4.1: blending empirical evidence with the deep-learning prior, rather than discarding the prior outright, preserves useful coverage in pocket regions the empirical fragment data does not sample.

### 6.5 Interpretability Analysis

Weight calibration (§4.2) yields a per-target weight vector that can be inspected directly. Table 4 reports the fitted weights for each target alongside PharmacoNet’s default weights. PharmacoNet uses a fixed set of interaction weights, heuristically defined from assumptions about the relative contributions of different chemical interactions across protein pockets. Because these fixed weights cannot account for variations in target-specific binding environments, we calibrate the interaction weights per target using Optuna, fit against each target’s own empirical fragment-docking data (§4.2). The resulting weights capture the target-specific interaction profile of each individual pocket, rather than the single population-averaged profile PharmacoNet otherwise assumes.

**Table 4.** MEL-Derived ScreenWeights vs. PharmacoNet default weights, per target.

| Interaction type | Default | CB2 | GPR91 | 5-HT<sub>2A</sub>R |
| --- | --- | --- | --- | --- |
| Hydrophobic | 1.0 | 1.00 | 1.00 | 1.00 |
| Aromatic | 4.0 | 2.03 | 2.78 | 1.14 |
| HBond_acceptor | 4.0 | 5.12 | 10.84 | 3.27 |
| HBond_donor | 4.0 | 12.91 | 3.25 | 9.17 |
| Halogen | 4.0 | 0.62 | 1.18 | 2.13 |
| Anion | 8.0 | 14.87 | 23.86 | 2.57 |
| Cation | 8.0 | 1.12 | 4.64 | 3.88 |

On GPR91, for instance, the fitted weights assign high importance to anionic and hydrogen-bond-acceptor interactions (23.86 and 10.84, respectively). This trend is consistent with the observed binding mode between GPR91 and its co-crystallized antagonist NF-56-EJ40 (Figure 4). The antagonist’s carboxylate group forms a salt bridge with protein residue R276 through the interactions, and its carbonyl group acts as a hydrogen-bond acceptor via a bridging water molecule. Thus, the fitted weights reflect the co-crystallized ligand’s binding mode, which capture the features of chemical interactions within the pocket of GPR91. See Supplementary Information §S6 for a caveat on interpreting weights fit from sparse pharmacophore evidence.

**Figure 4.**
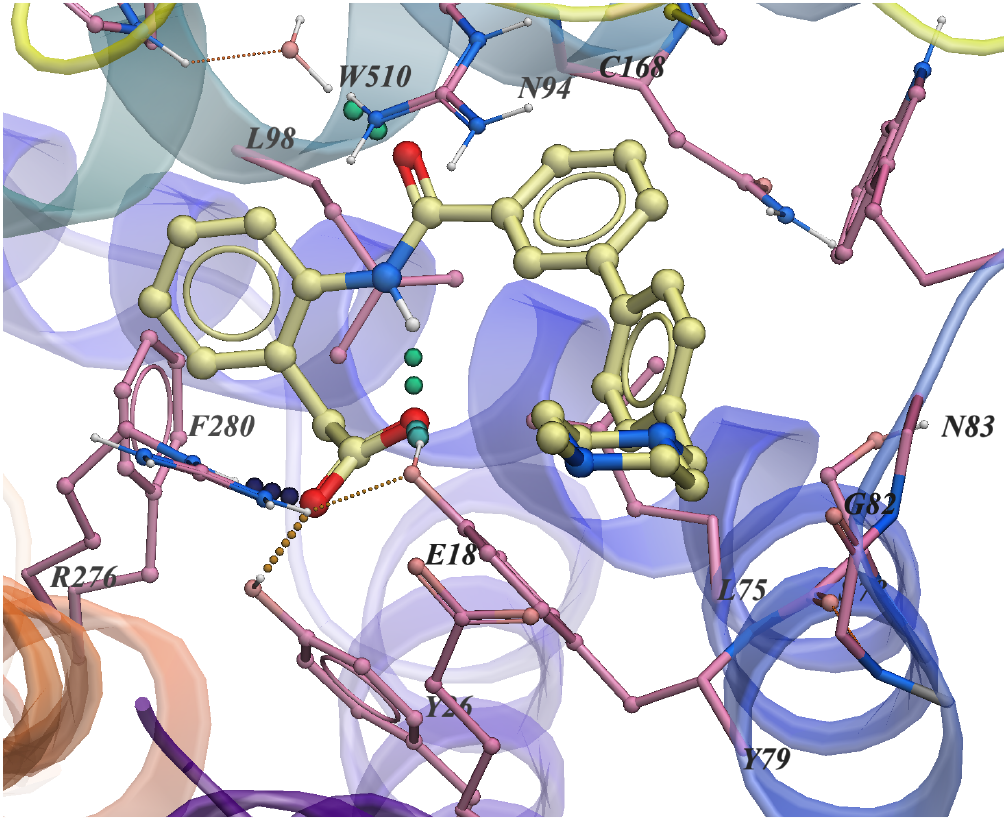
Co-crystallized binding mode of the GPR91 antagonist NF-56-EJ40 (PDB 6RNK).

## 7. Discussion & Conclusion

In this paper, we proposed MEL-Steered PharmacoNet, which specializes a general-purpose pharmacophore prescreening method to an individual target using signal the standard V-SYNTHES workflow already produces and normally discards, requiring no additional experimental data or retraining. Across three structurally distinct GPCR targets and both enrichment metrics, it consistently outperforms V-SYNTHES, vanilla PharmacoNet, DrugCLIP, and LigUnity. Ablations show geometry refinement and weight calibration correct distinct sources of error, each contributing independently and combining for further gains. Beyond enrichment, weight calibration yields a second, independent contribution: an interpretable, per-target account of which interactions the model relies on, giving practitioners a diagnostic view that PharmacoNet’s fixed weights cannot offer. While we instantiate this framework using PharmacoNet within V-SYNTHES, the underlying principle is modular and flexible: any screening pipeline that yields pocket-specific fragment docking evidence can provide the steering signal to fine-tune PharmacoNet. We leave this generalization, along with validation on non-GPCR targets, to future work.

## 8. Limitations & Ethical Considerations

We evaluate on three GPCR targets; generalization to other target classes (e.g., kinases, proteases, non-GPCR pockets) has not yet been tested. Our compound pools and docking scores depend on proprietary, licensed resources: the enumerated candidate and decoy sets are built from the Enamine REAL Space, a commercially licensed compound library, and docking is performed with ICM-Pro (Molsoft LLC), a licensed commercial docking engine. Both are widely used in industrial and academic drug discovery practice, but they limit exact reproducibility for researchers without access to these licenses. Like all other screening methods, our results are entirely docking-score based; we do not have experimental binding data to confirm that the enrichment gains reported in §6.3 translate into true wet-lab hit rates. MEL-Steered PharmacoNet’s ability to prioritize high-affinity binders is agnostic to therapeutic intent and could in principle be misused to design harmful compounds.

## GenAI Disclosure

Generative AI tools were used in a limited capacity. Claude was utilized for minor code construction, as well as for grammar and writing refinement.

## Acknowledgments

This work was supported in part by an NVIDIA Academic Grant, which provided GPU resources used in this study.

## Supplementary Information

**Table S1.** Pre-screening runtime for all baselines and MEL-Steered PharmacoNet, across three targets. Runtime in hours.

| Method | Runtime (h) ↓ |  |  |
| --- | --- | --- | --- |
|  | CB2 | GPR91 | 5-HT<sub>2A</sub>R |
| DrugCLIP | 22.82 | 17.02 | 21.78 |
| LigUnity | 1.72 | 1.58 | 1.65 |
| PharmacoNet | 2.22 | 2.31 | 2.73 |
| V-SYNTHES | 0.0 | 0.0 | 0.0 |
| MEL-Steered PharmacoNet | 1.24 | 1.19 | 1.16 |

### S1 Prescreening runtime comparison

In Table S1, the runtime we report and compare is restricted to the prescreening step alone: the computation each method uses to shortlist the top one million compounds from the target-specific compound space, rather than the pipeline’s total wall-clock time. The two remaining docking stages, MEL fragment docking and docking the final selected pool, are runtime-identical across all methods, so method-to-method comparison is only meaningful for the prescreening step between them. V-SYNTHES requires no prescreening computation at all: it selects the docking pool by fragment rank order directly, with no scoring pass over the enumerated compounds, so its prescreening runtime is effectively zero.

### S2 Embedding-based fast-screening methods

DrugCLIP and LigUnity represent a second class of generalist prescreening tools: both embed a protein pocket and a candidate ligand into a shared representation space and score candidates by embedding similarity rather than geometric matching [2, 3]. Like PharmacoNet, both are trained once across a broad population of protein– ligand interactions and applied in practice without target-specific adjustment. Both are trained and validated primarily against experimentally confirmed binding affinity rather than docking score, so ranking them by docking-score enrichment is not a strictly fair comparison; however, this embedding-based approach is nonetheless common practice as a prescreening layer in drug discovery, and we include both methods in our evaluation on that basis.

### S3 Weight Calibration Cross-Validation

Weights are selected via 5-fold cross-validation, stratified by docking-score percentile across six buckets (top 1%, 1–5%, 5–10%, 10–25%, 25–50%, 50–100%) to ensure each fold contains representative top-tier coverage. Cross-validation here serves as a diagnostic against search noise on a small population, to confirm the search is not simply chasing sampling noise in a single fixed fold assignment.

The deployed weight vector for each target comes from one further Optuna study, refit on the full population and warm-started by enqueueing the geometric mean of the five folds’ winning weight vectors as its first trial. We report the five folds’ held-out Spearman and NDCG@100 (mean ± std) as a cross-validated generalization estimate, and the final refit’s full-population score separately as an in-sample sanity check. The actual out-of-sample validation of the resulting weights is the enrichment factor evaluation on each target’s fully enumerated compound library (§6), obtained entirely outside this fitting procedure.

### S4 PharmacoNet Interaction-Type Weights

Each W_t_ in §4.2 is the scoring weight assigned to interaction type t : a per-type scaling factor applied to a matched pharmacophore hotspot, which determines how strongly a satisfied interaction of that type contributes to the overall graph-matching score S. PharmacoNet defines seven such types, each corresponding to a distinct class of non-covalent protein–ligand contact:

- Hydrophobic (default 1.0): nonpolar contacts between apolar ligand and pocket atoms, driven by the burial of nonpolar surface area.
- Aromatic (default 4.0): ring-stacking interactions (π–π) between an aromatic ligand ring and an aromatic pocket residue.
- Hydrogen-bond acceptor / donor (default 4.0 each): classical hydrogen bonds, in which the ligand presents an acceptor (lone pair) or donor (polarized H) group to a complementary donor or acceptor on the pocket.
- Halogen (default 4.0): halogen bonds, in which a halogen substituent (F, Cl, Br, I) on the ligand acts as an electrophile toward a pocket Lewis base.
- Cation / anion (default 8.0 each): electrostatic interactions, including salt bridges and cation–π contacts, between charged or partially charged ligand and pocket groups.

PharmacoNet’s default weights are fixed once across this typology and applied identically to every pocket; §4.2 recalibrates the six non-anchor weights per target using empirical MEL fragment-docking data.

### S5 Compound Filtering and Target-Specific Pool Construction

For each target, MEL fragments docked against the target pocket using ICM-Pro were retained by top docking score (top 1,000 fragments per target) and combinatorially expanded( each fragment’s capped position replaced by its full range of compatible synthons) into a target-specific candidate compound set. Enumerated compounds were filtered in two stages:

1. Physicochemical property filter: molecular weight ≤600 Da, cLogP ≤ 5.0, cLogS ≥ − 5.0, ≤5 hydrogen bond donors, ≤10 hydrogen bond acceptors, ≤10 rotatable bonds.
2. Substructure catalog filter: compounds matching PAINS, BRENK, NIH, or ZINC catalog entries were removed; surviving compounds were deduplicated by InChIKey.

This procedure yields 16,172,402 filtered compounds for CB2 (5ZTY), 14,928,903 for GPR91 (6RNK), and 16,111,451 for 5-HT_2A_R (7WC6). Each pool is constructed identically (same MEL library size, same top-1,000-fragment retention, same two-stage filtering) but is specific to its target, since fragment docking scores and therefore which fragments rank in the top 1,000 differ by target. These three pools constitute the fully enumerated compound space referenced in §6.2 and used as the common candidate pool for all prescreening methods and baselines evaluated in §6.

### S6 Weight Interpretability Caveat: Zero-Presence Interaction Types

The MEL-Derived ScreenWeights and one interpretability example are shown in Table 4 and §6.5. There is one important caveat for interpreting these weights: a weight is only meaningful if its corresponding interaction type is present in the pharmacophore model used for ScreenWeights fitting.

By Eq. (2), each weight W_t_ enters the PharmacoNet score S only through matched edges, (i → j), (i^′^→ j ^′^) where one of the two matched model nodes has type t . A target’s MEL-steered pharmacophore model (§4.1) can contain zero node of type t . This happens when no empirical contact of that type was found in the top-scoring MEL fragment population, and PharmacoNet’s deep-learning prior did not predict one either. When this is the case, W_t_ is never used in the score calculation for any candidate. The value reported in Table 4 for such a weight is whatever Optuna returns, and it should not be interpreted as reflecting the target’s interaction preferences.

Table S2 reports the number of matched nodes (n) of each interaction type in the MEL-steered pharmacophore model used to fit ScreenWeights (Table 4).

CB2’s pharmacophore model has no HBond_acceptor, HBond_donor, or Anion nodes; GPR91’s has no Aromatic nodes; and 5-HT_2A_R’s has no Anion nodes. Figure S1 confirms this pattern directly. The fitted weights at these five target-interaction type combinations, such as CB2’s Anion ScreenWeight of 14.87, carry no information about the target’s pocket preferences and should not be interpreted mechanistically. When interpreting Table 4, only ScreenWeights for interaction types with a nonzero number of matched nodes should be compared for a given target.

**Table S2.** Number of MEL-steered pharmacophore model nodes (n) per interaction type.

| Interaction type | CB2 | GPR91 | 5-HT<sub>2AR</sub> |
| --- | --- | --- | --- |
| Hydrophobic | 13 | 5 | 5 |
| Aromatic | 11 | 0 | 2 |
| HBond_acceptor | 0 | 1 | 1 |
| HBond_donor | 0 | 0 | 3 |
| Halogen | 1 | 4 | 8 |
| Anion | 0 | 1 | 0 |
| Cation | 4 | 1 | 1 |

**Figure S1.**
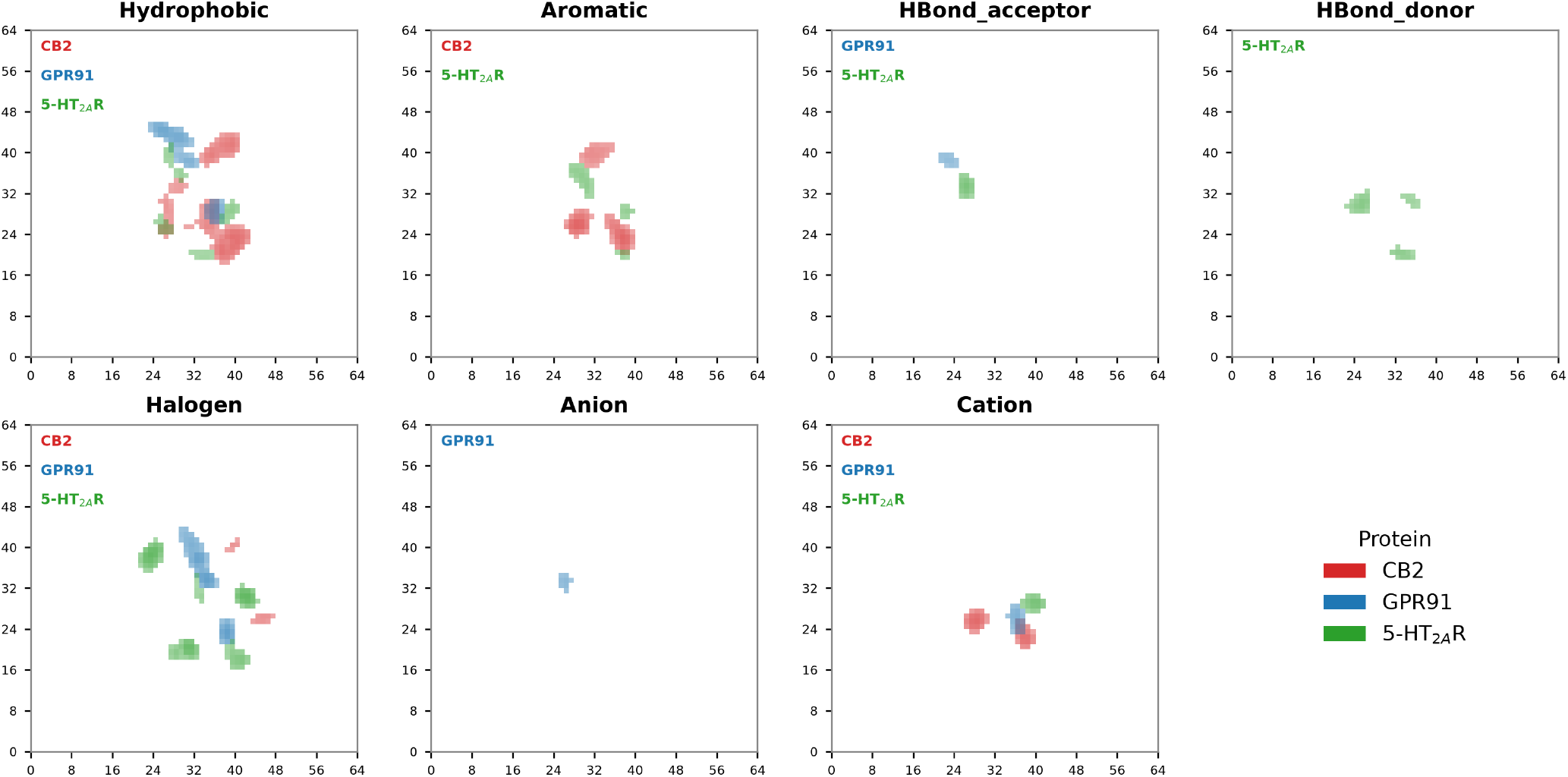
Per-target, per-interaction type coverage of the MEL-Steered pharmacophore models used to fit ScreenWeights (Table 4). Each panel shows the z-axis projection of matched pharmacophore nodes for one interaction type, colored by target (CB2, GPR91, 5-HT_2*A*_R). A target missing from a panel indicates zero matched nodes of that type.

## Footnotes

1 Pharmacophore-model and graph-matching illustrations adapted from [11].

2 Created in https://BioRender.com

